# Development of a ddPCR approach for the absolute quantification of soil microorganisms involved in atmospheric CO_2_ fixation

**DOI:** 10.1101/2024.03.25.586531

**Authors:** Marie Le Geay, Kyle Mayers, Martin Küttim, Béatrice Lauga, Vincent E.J. Jassey

**Author notes:** **Author contributions** MLG, BL and VEJJ conceived the study. VEJJ collected the samples in each site with the help of MK. MLG analyzed the samples with the help of BL and KM. MLG performed the statistical analyses. MLG wrote the first draft of the manuscript with the help of BL and VEJJ. All co-authors reviewed and contributed to the final form of the manuscript. **Data availability** Data related to this paper will be available from Figshare (10.xxxx/m9.figshare.c.xxxx) upon publication.

## Abstract

Carbon fixing microorganisms (CFMs) play a crucial role in soil carbon (C) cycling contributing to carbon uptake and sequestration through various metabolic pathways. Despite their significance, quantification of the absolute abundance of CFMs in soils remains elusive. This study employed a digital droplet PCR (ddPCR) approach to quantify the abundance of key and emerging CFM pathways in fen and bog across different depths (0-15 cm). Targeting total prokaryotes (*16S rRNA* gene), oxygenic phototrophs (*23S rRNA* gene), aerobic anoxygenic phototrophic bacteria (AAnPB, *pufM* gene), and chemoautotrophs (*cbbL* gene), we optimized ddPCR conditions to achieve absolute quantification of these genes. Overall, our results revealed that oxygenic phototrophs were the most abundant CFMs, constituting 12% of total prokaryotic abundance, followed by chemoautotrophs (10%) and AAnPBs (9%). Fen exhibited higher gene concentrations than bog. Depth variations were also observed, differing between fen and bog for all genes. Our findings highlight the abundance of oxygenic phototrophs and chemoautotrophs in peatlands, challenging previous estimations that relied solely on oxygenic phototrophs for microbial CO_2_ fixation assessments. Incorporating absolute gene quantification is crucial for a comprehensive understanding of microbial contributions to soil processes, shedding light on the intricate mechanisms of soil functioning in peatlands.

## Introduction

Carbon fixing microorganisms (CFMs) are key drivers of the carbon (C) cycle as they assimilate atmospheric CO_2_ and contribute to the soil organic C sequestration (Yuan et al., 2012; Ge et al., 2013; Liu et al., 2018). CFMs fix CO_2_ through six major metabolic pathways, namely, the reductive pentose phosphate cycle (also known as Calvin-Benson-Bassham, CBB cycle), the reductive citrate cycle (rTCA cycle), the 3-hydroxypropionate bi-cycle (3-HP cycle), the 3-hydroxypropionate/4-hydroxybutyrate cycle (3-HP/4-HB cycle), the dicarboxylate-hydroxybutyrate cycle (DC/4-HB cycle) and the reductive acetyl-CoA pathway (also known as Wood-Ljungdahl pathway) (Liu et al., 2018; Huang et al., 2022). The CBB cycle is the predominant pathway utilized by microorganisms in soils (Bay et al., 2021). It is present across different taxonomic levels (bacteria and protists), notably in green algae, diatoms, cyanobacteria and aerobic eubacteria (Berg, 2011; Yuan et al., 2012). Oxygenic phototrophs and chemoautotrophs are two major CFMs using the CBB cycle to fix atmospheric CO_2_. Oxygenic phototrophs derive their energy from light to assimilate CO_2_ through photosynthesis, whereas chemoautotrophs use reduced chemical compounds such as sulfur compounds, molecular hydrogen and reduced metals to assimilate CO_2_ (Hügler and Sievert, 2011). Potential CO_2_ fixation through the CBB cycle also recently emerged in aerobic anoxygenic phototrophic bacteria (AAnPBs). Recent studies found AAnPB strains possessing (and expressing) genes of the CBB cycle (Graham et al., 2018; Tang et al., 2021). Even though CFMs are now well recognized for their role in autotrophic CO_2_ fixation in soils, their absolute quantification remains elusive (Liao et al., 2023). As microbial CO_2_ assimilation rates are closely linked to CFMs abundance (Hamard et al., 2021a, 2021b; Liao et al., 2023), it is essential to provide absolute quantification of these microorganisms in soils to better assess their contribution in C fixation in terrestrial ecosystems.

Absolute quantification of microorganisms can be laborious notably in complex matrices such as soils. Numerous approaches are available with their own strengths and flaws (Xiaofan Wang et al., 2021). Methods based on cells counts such as fluorescence microscopy or flow cytometry have been widely used (Bressan et al., 2015; Frossard et al., 2016), as have methods based on adenosine triphosphate quantification (Karl, 1980; Hammes et al., 2010) or phospholipids fatty acids analysis (PLFA, Frostegård and Bååth, 1996). However, these methods (except PLFA) do not allow differentiation between different bacterial groups. Since the development of Polymerase Chain Reaction (PCR, Mullis et al., 1986), different molecular approaches, such as real-time PCR (qPCR, Higuchi et al., 1993), digital PCR (dPCR, Vogelstein and Kinzler, 1999) and digital droplet PCR (ddPCR, Hindson et al., 2011) have emerged to count either total bacteria (using primers targeting the *16S rRNA* gene) or specific populations of bacteria and their associated functions (using primers targeting specific genes; Fig. S1). qPCR is based on the monitoring of the amplification after each PCR cycle by using a fluorescent probe (Higuchi et al., 1993; Pinheiro et al., 2012; Hou et al., 2023). However, this method requires external calibration and normalization with endogenous controls (Hindson et al., 2011). It is also sensitive to inhibitors and to the initial amount of the target since low concentrations will hardly be detected (Hindson et al., 2011; Pinheiro et al., 2012; Taylor et al., 2017; Hou et al., 2023). On the other hand, ddPCR does not require external calibration or normalization. This method uses water-in-oil droplets allowing single molecule amplification. Droplets can either contain the target molecule or not. After being amplified with a fluorescent probe or labeling dye, it is possible to discriminate between positive and negative droplets (see Hindson et al., 2011). Several studies have compared qPCR and ddPCR and found that ddPCR had better precision, repeatability, sensitivity and stability than qPCR (Zhao et al., 2016; Taylor et al., 2017; Xue et al., 2018; Wang et al., 2022). Nowadays, ddPCR is increasingly used in environmental studies (see Hou et al., 2023 for a complete review), even though several challenges are faced and require appropriate optimization of ddPCR parameters (see Kokkoris et al., 2021). Adjusting many factors relating to the PCR reaction, including concentration of template DNA, thermocycling conditions, and the threshold setting used to discriminate positive and negative droplets can notably influence the detection of target DNA from environmental samples (Witte et al., 2016; Rowlands et al., 2019; Kokkoris et al., 2021).

In this study, we aim to provide an absolute quantification of the main (oxygenic phototrophs and chemoautotrophs) and emerging (AAnPBs) CFMs in peatlands. As peatlands are a major C reservoir (Nichols and Peteet, 2019), quantifying the main CFMs will enhance our understanding of peatland C cycling. More precisely, we aimed to (1) optimize ddPCR conditions to enumerate total prokaryotes and specific groups of CFMs (oxygenic phototrophs, chemoautotrophs and AAnPBs), (2) apply optimized ddPCR to target CFMs in two peatland types (moderately rich fen and open bog) and (3) at different depths (0-15 cm). We hypothesized that oxygenic phototrophs, who can be found within bacteria and protists, will be the most abundant CFMs in peatlands. We also forecast that CFMs absolute abundance will differ between the bog and the fen as these two peatland habitats harbor different biotic and abiotic factors. We further hypothesized that oxygenic phototrophs and AAnPBs will be less abundant with depth as the amount of light decreases, thus favoring chemoautotrophs.

## Experimental procedure

### Sampling and DNA extraction

Peat samples were collected in two peatlands, Counozouls (France) and Männikjärve (Estonia). Counozouls mire is located in the French Pyrenees mountains (42°41’16N” - 2°14’18”E – 1.374 m a.s.l). It is a moderately rich fen belonging to the Special Area of the Natura 2000 Conservation site “Massif du Madres Coronat”. The peatland of Männikjärve is an open bog located in the Endla mire system in Central Estonia (58°52’26.4”N - 26°15’03.6”E – 82 m a.s.l). At each site, we collected three cores in similar and homogeneous habitat by cutting peat soil. Then, the cores were cut into three depths corresponding to the living layer (D1; 0-5 cm), the decaying layer (D2; 5-10 cm), and the dead layer (D3; 10-15 cm). At each depth, few grams of peat were collected, cut in small pieces, homogenized and placed in sterile 5 mL Eppendorf tubes containing 3 mL of RNA*later* (ThermoFisher). The tubes were stored at -20°C upon our arrival in the laboratory. DNA was extracted using the DNeasy PowerSoil Pro Kit (Qiagen) following manufacturer’s instructions. After elution of DNA (70 µL in final solution), DNA concentration was quantified using a Nanodrop ND-1000 spectrophotometer. Extracts were then stored at -20°C prior to DNA amplification.

### Primer choice and design

Total concentration of prokaryotes was obtained by targeting the *16S rRNA* gene. Microorganisms involved in oxygenic photosynthesis were quantified using the *23S rRNA* gene while chemoautotrophs were quantified using the *cbbL* gene that encodes the large subunit of RubisCO form IA (Kusian and Bowien, 1997; Alfreider and Bogensperger, 2018). We used the *pufM* gene that encodes for the M subunit of type II photochemical reaction center to target AAnPBs (Achenbach et al., 2001; Béjà et al., 2002). To select the primers, we first searched the literature and identified primer pairs that have been already used either in ddPCR or in qPCR. We used the primer pairs L/Prba338f and K/Prun518r for prokaryotes (Øvreås et al., 1997), cbbLR1F/cbbLR1inR for chemoautotrophs (Selesi et al., 2007) and pufMforward557/pufMreverse750 for AAnPBs (Table 1; Du et al., 2006). However, we could not find suitable primers matching a short region of the *23S rRNA* gene targeting oxygenic phototrophs with satisfying length and degeneracies. Therefore, we designed a new set of primers for ddPCR.

**Table 1.**
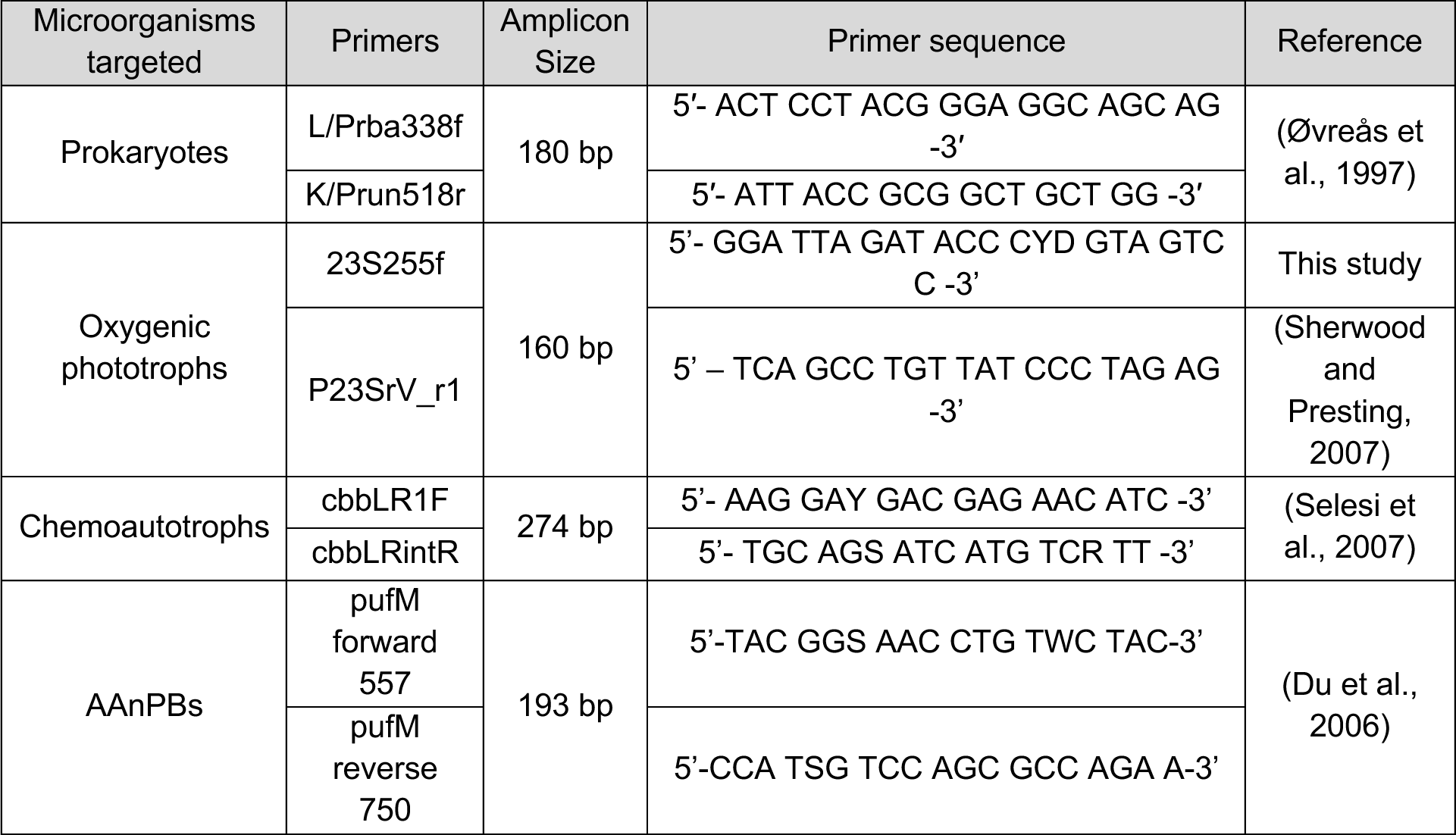
Primer pairs used in this study.

To do so, we first conducted a PCR on peat samples to target the *23S rRNA* gene using the existing primers P23SrV-f1 with P23SRV-r1 (Sherwood and Presting, 2007) and sequenced PCR amplicons. The PCR program was run in a total volume of 50 μL containing 25 μL of AmpliTaq GoldTM Master Mix (applied biosystem, ThermoFisher), 19 μL of ultrapure water, 1 μL of forward and 1 μL of reverse primer (final concentration of 20 μM) and 2 μL of DNA template. The PCR reaction conditions were an initialization of 10 min at 95°C followed by 35 cycles of 1 min at 94°C, 45 s at 55°C, 45 s at 72°C and a final extension step of 10 min at 72°C. PCR quality was assessed using 1.65% agarose gel electrophoresis. The high throughput sequencing was performed by the GeT-PlaGe plateform (Genotoul, Toulouse, France) using Illumina MiSeq v3 technology. Based on the results of the sequencing we designed the new forward primer 23S255f to target a short region of the *23S rRNA* gene. We aligned all the sequences obtained from sequencing using Clustal Omega (EMBL-EBI) and blasted (nBLAST with NCBI) these sequences to ensure that they corresponded to the *23S rRNA* gene. Then, using BioEdit v5.0.9 we generated a consensus sequence based on the aligned sequences (Fig. S2). We searched in this consensus sequence a conserved region and checked every degenerated base to create a manually curated sequence. We examined if this sequence matched guidelines for q/dPCR (Rodríguez et al., 2015) (amplicon length <250 pb, GC content of 50-60%, annealing temperature of 50-65°C, no secondary structure and primer-dimer, no repetition of Gs and Cs longer than 3 bases, Table S1) and finally obtained the new forward primer 23S255f: 5’-GGA TTA GAT ACC CYD GTA GTC C -3’. By using the new primer set, the forward primer 23S255f with the reverse primer P23SRV-r1 (∼160 pb), we targeted oxygenic phototrophs.

### ddPCR conditions

Absolute abundance of genes was measured using digital droplet PCR (ddPCR, BioRad). The ddPCR reactions were run in a total volume of 20 µL on a DX200 instrument (BioRad) with 10 µL of EvaGreen Supermix (BioRad, 1X), 0.5 or 0.3 µL of each primer (final concentration 250 nM or 150 nM respectively) and 4 µL of ultrapure water. Template DNA (5 µL) diluted at 10, 100 or 1,000 times was added to the reaction mix. This mixture was then emulsified with QX200 Droplet Generation Oil for EvaGreen (BioRad) using the QX200 Droplet Generator (BioRad) and was manually transferred into a 96-well PCR plate. The plate was heat-sealed with a foil seal and then placed on a C1000 Touch Thermocycler with deep-well module (BioRad) to run the PCR using different programs (see section *‘ddPCR optimization’*). Following amplification, plates were equilibrated for at least 10 min at room temperature. Then, plates were loaded on a QX200 Droplet Reader (BioRad) and the fluorescence was read and analyzed using QuantaSoft software. Threshold for positive and negative droplets were manually defined using ultrapure water as a negative control and DNA extracted from different cultures of microorganisms as positive and negative controls. *Escherichia coli* DNA was used as a positive control for prokaryotes and as a negative control for the other genes. *Micromonas pusilla* (a micro-algae) DNA was used as a positive control for oxygenic phototrophs. QuantaSoft provides a final concentration of target copies.µL^-1^ of ddPCR reaction. We first calculated the total concentration in 20 µL of ddPCR reaction and normalized it by the weight of dry peat to obtain a final concentration in target copies.g^-1^ of dry peat (copies.g^-1^ DW).

### ddPCR optimization

For all the assays we tested two primer concentrations (150 nM and 250 nM); a temperature gradient based on the theorical annealing temperature of the primers; different dilution of the template (1/10, 1/100 and 1/1,000) and different species DNA to find controls (*E. coli*, *M. pusilla*). We defined thermocycling conditions as 98°C for 5 min followed by 40 cycles of 94°C for 30 s, different temperature or gradient of temperature for 30 s, followed by 5 min at 4°C and 98°C for 10 min. The ramp rate was set up at 2°C/s. When required, these parameters were adjusted to optimize the assays. For two assays (*16S* and *23S rRNA* genes) we used the following PCR conditions: 98°C for 5 min followed by 40 cycles of 94°C for 30 s, 61°C and 57.6°C for 1 min, followed by 5 minutes at 4°C and 98°C for 10 min with a ramp rate stetted up at 2°C/s and for the others (*cbbL* and *pufM* genes) we used: 98°C for 5 min followed by 45 cycles of 94°C for 1 min, 53°C and 50.2°C for 2 min, followed by 5 min at 4°C and 98°C for 10 min with a ramp rate stetted up at 1°C/s (Table 2).

**Table 2.** Optimized ddPCR parameters including primers concentration, DNA template dilution, optimal annealing temperature, ddPCR thermocycling conditions and positive and negative controls, UPW = ultra-pure water

| Primers | L/Prba338f with K/Prun518r | 23S255f/ P23Srv-r1 | cbbLR1F/ cbbLRintR1 | pufMfwd557/ pufMrev750 |
| --- | --- | --- | --- | --- |
| Gene targeted | 16S *rRNA* | 23S *rRNA* | *cbbL* | *pufM* |
| Primers concentration | 150 nM | 250 nM | 250 nM | 250 nM |
| DNA dilution | 1/1,000 | 1/100 (D1, D2) and 1/10 (D3) | 1/10 | 1/100 (D1, D2) and 1/10 (D3) |
| Annealing temperature | 61 °C | 57.6 °C | 53 °C | 50.2 °C |
| ddPCR conditions | 98 °C – 5 min<br>40 cycles :<br>94 °C – 30 s<br>61 °C – 1 min | 98 °C – 5 min<br>40 cycles :<br>94 °C – 30 s<br>57.6 °C – 1 min | 98 °C – 5 min<br>45 cycles :<br>94 °C – 1 min<br>53 °C – 2 min | 98 °C – 5 min<br>45 cycles :<br>94 °C – 1 min<br>50.2 °C – 2 min |
|  | 4°C – 5 min<br>98°C – 10 min<br>Ramp rate =<br>2°C/s | 4°C – 5 min<br>98°C – 10 min<br>Ramp rate =<br>2°C/s | 4°C – 5 min<br>98°C – 10 min<br>Ramp rate = 1°C/s | 4°C – 5 min<br>98°C – 10 min<br>Ramp rate = 1°C/s |
| Positive control | *E. coli* DNA | *M. pusilla* DNA | / | / |
| Negative control | UPW | *E. coli* DNA<br>UPW | *E. coli* DNA<br>UPW | *E. coli* DNA<br>UPW |

### Statistical analysis

Data were visually checked and tested for normality and homoscedasticity, and log-transformed when necessary. Normality was assessed using Shapiro-Wilk test (Shapiro and Wilk, 1965). In order to see if the genes concentrations differed according to location or depth we conducted a one-way ANOVA. We also tested if the concentration significantly differed according to the gene targeted. Post-hoc test (Tukey honestly significant difference test) was further applied when a significant difference was found between location, depth and genes (Keselman and Rogan, 1977). Statistical analyses were performed using RStudio v12.0 (RStudio Team, 2020) and graphical representations were done using ggplot2 v3.4.0 (Wickham, 2016).

## Results and Discussion

### Optimized ddPCR succeed to amplify universal and specific genes from peat samples

We tested different ddPCR conditions to obtain both a good amplification of the target genes and a separation between positive and negative clouds of droplets (Fig. 1). When numerous droplets were present without a clear separation (rain), ddPCR conditions were optimized (Fig. S3-S6) following recommendations from previous studies (Witte et al., 2016; Rowlands et al., 2019; Kokkoris et al., 2021). We notably optimized the primers annealing temperature (better separation of the droplets), the dilution of the target (dilution of potential inhibitors and avoid saturation of the target, Fig. S3 well A05), the thermocycling conditions (denaturation/annealing/elongation times, number of cycle and ramp rate leading to compacts clouds of positive and negative droplets well separated) and we used positive and negative controls to set an appropriate thresholds (Witte et al., 2016; Rowlands et al., 2019; Demeke et al., 2021; Kokkoris et al., 2021). Results of the optimization are presented in Table 2 and Supplementary Figures S3, S4, S5 and S6.

**Figure 1.**
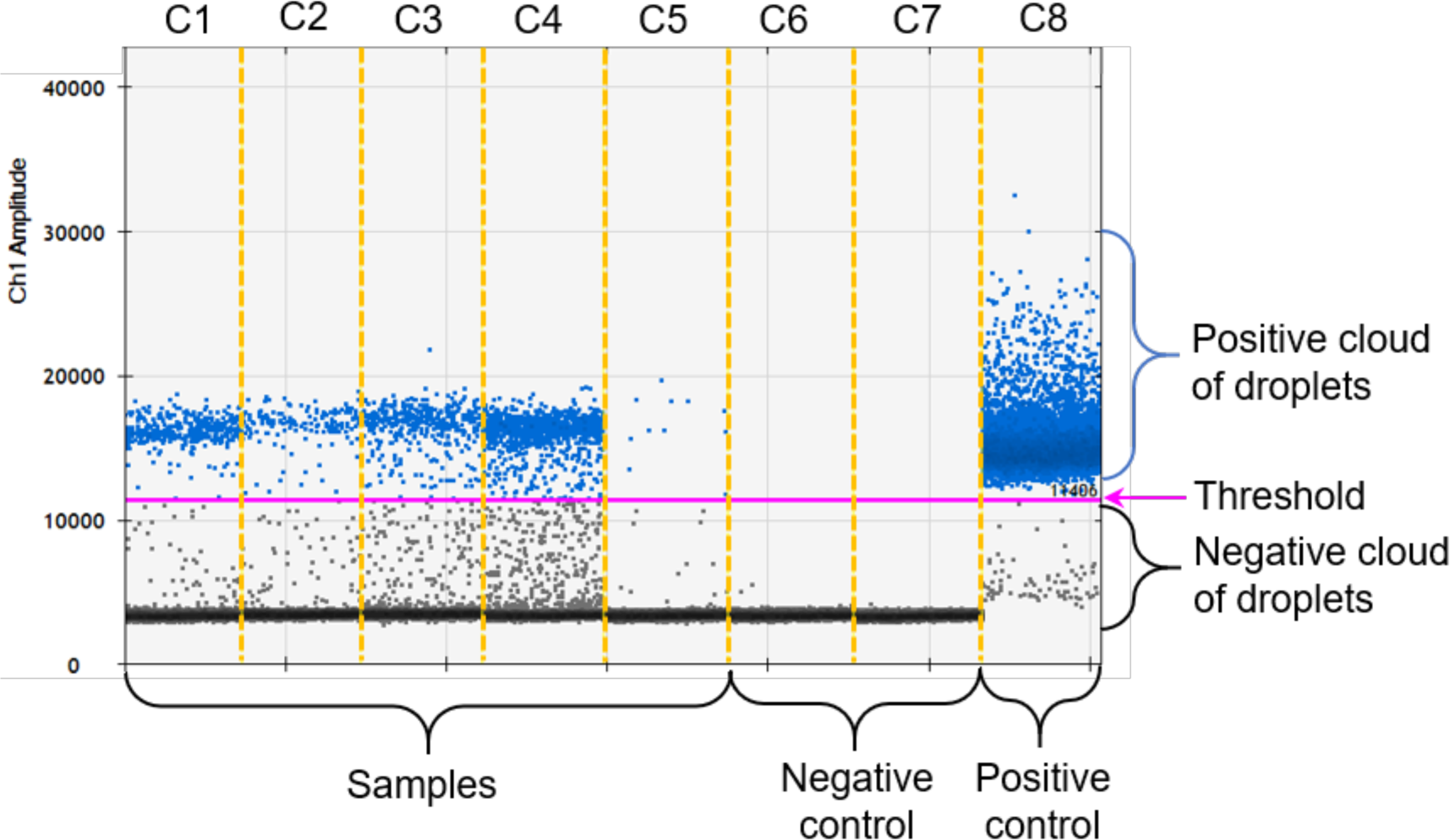
Example of results of ddPCR obtained for the *23S rRNA* gene after analysis of raw data with QuantaSoft software. C1 to C5 = soil samples, C6 = ultrapure water (negative control), C7 = *E. coli* DNA (corresponding here to a negative control) and C8 = *M. pusilla* DNA (positive control). Horizontal purple line corresponds to the threshold manually set thanks to negative and positive controls. Blue dots correspond to positive droplets (containing target DNA) and black dots to negative droplets (not containing target DNA).

Once the assays were otpimised, we used the ddPCR parameters defined to amplify DNA from samples of the two peatlands location (Table 2). We retrieved *16S rRNA* gene concentrations of 10.3 ± 2.76 x 10^6^ copies.g^-1^ DW in Counozouls and 2.11 ± 0.21 x 10^6^ copies.g^-1^ DW in Männikjärve. Previous peatlands studies found total bacterial concentrations from 1 x 10^6^ copies.g^-1^ DW or copies.mL^-1^ (Gilbert et al., 1998; Lew et al., 2016) to 1 x 10^10^ copies.g^-1^ DW or copies.mL^-1^ (Lin et al., 2012; Wen et al., 2018) which is consistent with our findings. Our data further showed that CFMs genes concentrations were very similar (*P* > 0.05, ANOVAs; Fig. 2). Average *23S rRNA* gene concentration was 14.0 ± 2.62 x 10^5^ copies.g^-1^ DW in Counozouls and 6.98 ± 1.67 x 10^5^ copies.g^-1^ DW in Männikjärve. Again, this is consistent with other findings from soils (Jassey et al., 2022) and in particular peatlands where concentrations around 1 x 10^5^ – 10^-6^ copies.g^-1^ DW have been found using microscopy and qPCR (Gilbert et al., 1998; Jassey et al., 2015; Reczuga et al., 2018; Tang et al., 2018). On average, *cbbL* gene concentration was 9.90 ± 2.36 x 10^5^ copies.g^-1^ DW in Counozouls and 2.57 ± 0.43 x 10^5^ copies.g^-1^ DW in Männikjärve. Similar abundances were retrieved in soils using qPCR (Xiao et al., 2014; Keshri et al., 2015; Xiayu Wang et al., 2021; Yin et al., 2022). For the *pufM* gene, 8.17 ± 1.59 x 10^5^ copies.g^-1^ DW were retrieved in Counozouls and 3.24 ± 0.61 x 10^5^ copies.g^-1^ DW in Männikjärve. Again, these concentrations were comparable to AAnPBs concentrations found in soils (1-50 x 10^5^ copies.g^-1^) or in aquatic environments (6 x 10^4^ - 12 x 10^5^ copies.mL^-1^) (Lew et al., 2016; Sato-Takabe et al., 2016, 2020; Tang et al., 2018). All together, these results showed that ddPCR is a suitable and reliable method to quantify absolute gene concentrations from complex soil matrices, like peat. Indeed, peat samples are usually rich in humic substances that are hardly removed during DNA extraction (Delarue et al., 2011). Their presence in DNA sample can inhibit Taq DNA polymerase and the fluorescence signal of double-stranded DNA binding dyes (Sidstedt et al., 2015). By optimizing ddPCR parameters we were able to overcome this issue and improve the separation between positive and negative droplets.

**Figure 2.**
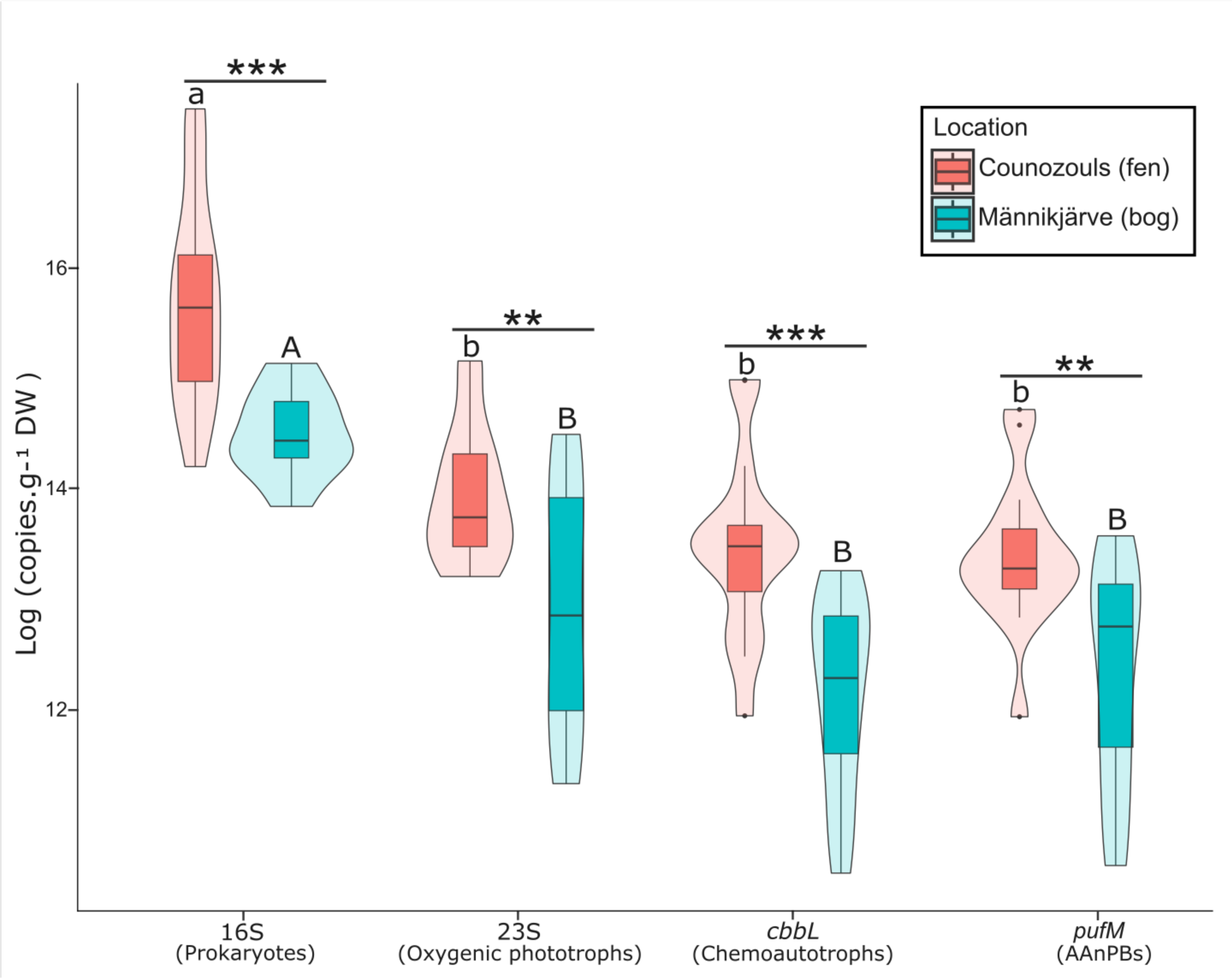
Absolute quantification of *16S rRNA* gene targeting prokaryotes, *23S rRNA* gene targeting oxygenic phototrophs, *cbbL* targeting chemoautotrophs and *pufM* targeting AAnPBs (all depth confounds) in Counozouls (fen) and Männikjärve (bog). Boxplots represent the logarithm of the total genes’ copies.g^-1^ DW and violin plots show the shape of data distribution. The horizontal black line in the box corresponds to the median and the edges of the box to 1^st^ and 3^rd^ quartiles. The vertical black line on the top and bottom of the boxplot represent respectively the highest and lowest values in the ± 1.5 interquartile range. Black dots correspond to the outliers. Lowercase and uppercase letters represent the difference between genes in Counozouls and Männikjärve, respectively. Horizontal bars with * represent the difference between peatlands (Counozouls and Männikjärve) for each gene, ** = *p* < 0.01; *** = *p* < 0.001. n = 15 for each gene at each location.

### The abundance of CFMs differ between fen and bog and with depth

Overall, we found that gene concentrations in the fen were up to 4 times greater than gene concentrations in the bog (*P* < 0.05 for all genes, ANOVAs; Fig. 2). This pattern agrees with previous findings on total bacteria (Lin et al., 2012; Mpamah et al., 2017; Xu et al., 2021) and photoautotrophic bacteria (Hamard et al., 2021a; Sytiuk et al., 2022) abundances assessed by either inverted microscopy, flow cytometry or PLFA assays. As a moderately rich fen and an open bog, Counozouls and Männikjärve present different nutrient contents, acidity and vegetation cover (Sytiuk et al., 2022) suggesting that these conditions are strong determinants of CFM abundance in peatlands. This finding corroborates previous findings on microbial CO_2_ fixation in soils, where nutrient and water content (Guo et al., 2015; Zhao et al., 2021), soil pH and soil organic carbon (SOC) (Lin et al., 2012; Nowak et al., 2015; Lew et al., 2016; Zhao et al., 2021) have been shown to influence microbial abundance in general and microbial CO_2_ fixation in particular. Additionally, or alternatively, our data also suggest that climate may determine CFM abundances. Counozouls and Männikjärve are located at different latitudes, and thus harbor different climate with warmer and wetter conditions in Counozouls than in Männikjärve (Sytiuk et al., 2022). Thus, warmer and wetter conditions in Counozouls may have increased the abundance of CFMs as previously shown for peatlands microbes (Le Geay et al., 2024), but also for microbial biomass in soils (Yuan et al., 2012; Xu et al., 2013). Although the exact environmental drivers of CFM abundance in peatlands still need to be identified, our results suggest that complex interactions between climate, peat properties and vegetation cover might determine CFM abundance in peatlands. Alternatively, DNA found in peat can be a mix of DNA from active cells, dead cells or even extracellular DNA (Pearman et al., 2022), which could explain the observed high abundance in the fen. Physicochemical conditions found in Counozouls may better preserve the DNA letting both active and dead cells preserved. More site comparisons will be required in the future to further understand this observed pattern.

In addition to peatland type effect, our data evidenced a net depth effect but with opposite patterns between the fen and the bog (Fig. 3). In Counozouls, all gene concentrations aside from *23S rRNA* and *pufM* increased (*P* < 0.05, ANOVAs; Fig. 3; Tables S2-S5) while in Männikjärve, all gene concentrations but *16S rRNA* decreased with depth (*P* < 0.05, ANOVAs; Fig. 3; Tables S2-S5). More particularly, we found that the concentration of the *23S rRNA* gene did not vary with depth ( *P* > 0.05) except between D2 and D3 samples in Counozouls (*P* < 0.05; Fig. 3; Table S3), while it gradually decreased with depth in Männikjärve (*P* < 0.001; Fig. 3; Table S3). As oxygenic phototrophs biomass is tightly linked to light availability (Bengtsson et al., 2018), we were expecting such a decrease of the *23S rRNA* gene with depth. We tentatively explain the high abundance of the *23S rRNA* gene with depth in Counozouls by the dominance of cyanobacteria in the oxygenic phototrophic community (Fig. S7). Indeed, cyanobacteria can capture light at low intensities thanks to highly adaptable eco-physiological traits (Carey et al., 2012) and could therefore remain abundant with depths in Counozouls.

**Figure 3.**
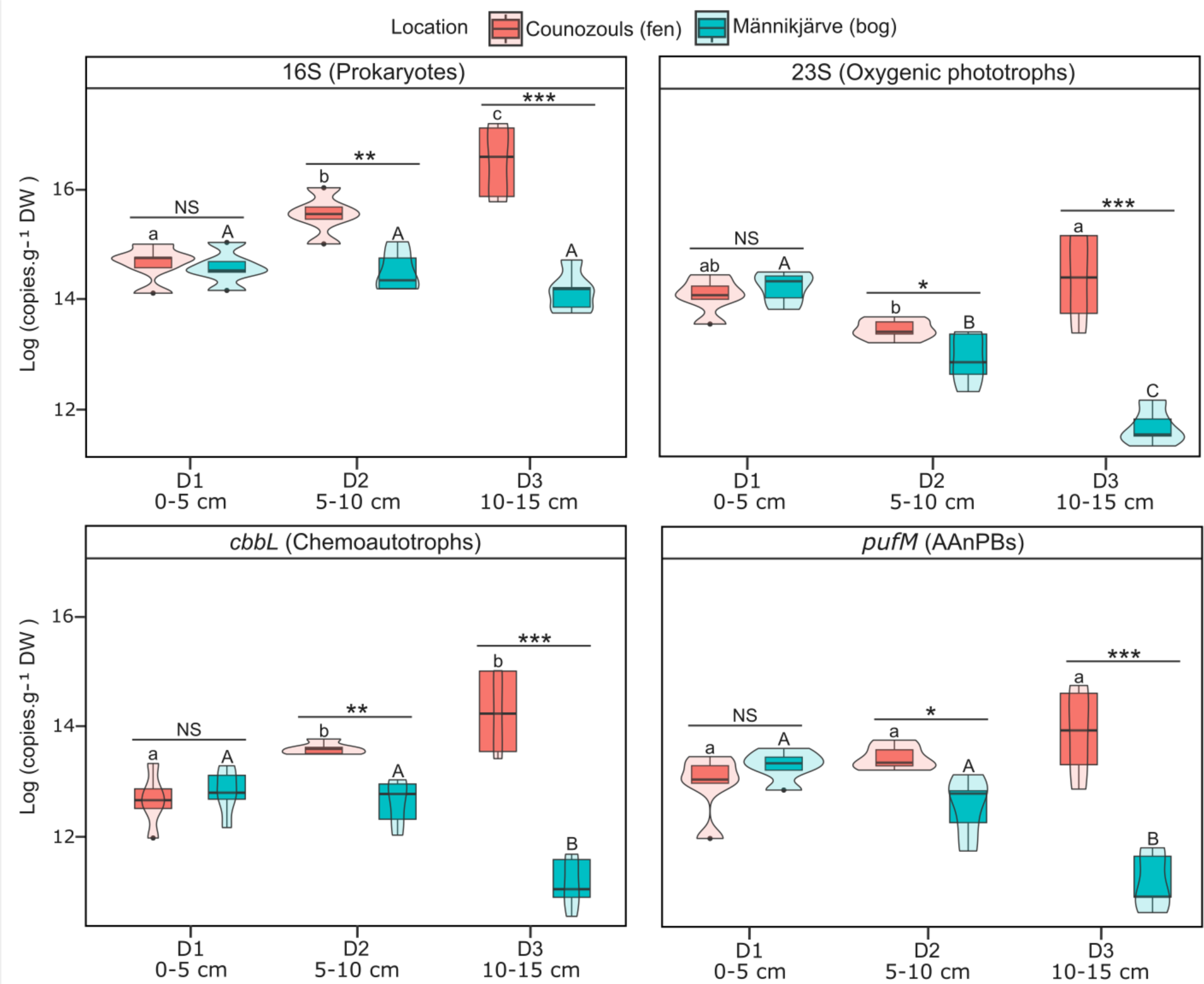
Absolute quantification of *16S rRNA* gene targeting prokaryotes, *23S rRNA* gene targeting oxygenic phototrophs, *cbbL* targeting chemoautotrophs and *pufM* targeting AAnPBs at different depths in Counozouls (fen) and Männikjärve (bog). Boxplots represent the logarithm of the total genes’ copies.g^-1^ DW and violin plots show the shape of data distribution. The black line in the box corresponds to the median and the edges of the box to 1^st^ and 3^rd^ quartiles. The vertical black line on the top and bottom of the boxplot represent respectively the highest and lowest values in the ± 1.5 interquartile range. Black dots correspond to the outliers. Lowercase and uppercase letters represent the difference between genes in Counozouls and Männikjärve, respectively. Horizontal bars with * represent the difference between peatlands (Counozouls and Männikjärve) at each depth. NS = not significant (*p* > 0.05); * = *p* < 0.05; ** = *p* < 0.01; *** = *p* < 0.001. n = 5 for each microbial group at each depth for each location. D1 (0-5 cm) corresponds to the living layer, D2 (5-10 cm) to the decaying layer and D3 (10-15 cm) to the dead layer.

Concentration of the *cbbL* gene targeting chemoautotrophs significantly increased with depth in Counozouls (*P* < 0.05, ANOVA) whereas it significantly decreased with depth in Männikjärve (*P* < 0.001; ANOVA; Fig. 3; Table S4). Chemoautotrophs are notably studied in deep sea vents, coastal sediments, and soils, where reduced compounds such as sulfur compounds, molecular hydrogen and reduced metals are particularly abundant and facilitate chemoautotrophy (Nakagawa and Takai, 2008; Boschker et al., 2014; Gomez-Saez et al., 2017; Yang et al., 2017; Wang et al., 2023). Usually chemoautotrophs are active in the first centimeters of coastal sediments as conditions are limiting in the deeper layers. We retrieved such a pattern in Männikjärve but not in Counozouls, suggesting that geochemical conditions for chemoautotrophs might be less limiting in Counozouls. For the *pufM* gene that targets AAnPBs, we did not see a significant difference in gene concentrations between depth in Counozouls (*P* > 0.05). In Männikjärve we found a significant decrease of the *pufM* gene concentration with depth (*P* < 0.001; ANOVA; Fig. 3; Table S5). Being obligate aerobic microorganisms, AAnPBs have mainly been found in the euphotic zone of aquatic systems (Koblížek et al., 2003). They are often described as highly active part of microbial communities; however, we are lacking information regarding their importance in soil communities as well as their metabolic capacities with only a few studies working on AAnPBs in soils (Feng et al., 2011; Tang et al., 2018). Our data show that AAnPBs are more abundant in fens than in bogs, suggesting they could play an important ecological role in these ecosystems.

### Ratios of oxygenic phototrophs, chemoautotrophs and AAnPBs compared to total prokaryotes reveals their equally importance in peatlands

Our data revealed that compared to total prokaryotes, prokaryotic oxygenic phototrophs, chemoautotrophs and AAnPBs were equally abundant. Prokaryotic oxygenic phototrophs and chemoautotrophs represented ∼8% and ∼10% of total prokaryotes in Counozouls and ∼16% and ∼10% in Männikjärve. These results showed that chemoautotrophs were more or less as abundant as oxygenic phototrophs, and possibly equally contributed to CO_2_ fixation in peatlands. This contradicts the first estimates of phototrophic and chemoautotrophic CO_2_ fixation rates in peatlands showing that chemoautotrophic CO_2_ assimilation represented less than 1% of oxygenic phototrophic CO_2_ fixation (Gilbert et al. 1998). Further work quantifying chemoautotrophic CO_2_ fixation in peatlands will be necessary in the future. Also, the use of RNA quantification instead of DNA may better evidence abundance patterns between phototrophs and chemoautotrophs. Furthermore, our results show that AAnPBs represented ∼8% of total prokaryotes in Counozouls and ∼15% of total prokaryotes in Männikjärve, suggesting they may play a significant role in the C cycle of peatlands as some strains possessing CBB cycle genes can actively fix atmospheric CO_2_ (Graham et al., 2018; Tang et al., 2021). Altogether these results show that further work is urgently needed to better quantify the contribution of microorganisms to peatland CO_2_ uptake as current estimates based on sole oxygenic phototrophs (Hamard et al., 2021a) may be strongly underestimated. Soils are also facing important climate and land use changes which can profoundly affect soil microbiomes and their functions (Solomon et al., 2007; Lode et al., 2017; IPCC, 2023). Changing soil microbiome can notably impact C fixation and emission (Blankinship et al., 2011; Zhou et al., 2020; Li et al., 2022; Le Geay et al., 2024). These changes could lead to enhanced microbial CO_2_ fixation in some soils, which could mitigate global changes (Qiu et al., 2020; Strack et al., 2022; Le Geay et al., 2024). Therefore, knowing the contribution of the different CFMs and their metabolic pathway to soil microbial CO_2_ fixation is also a key forecast about the changes in soils C cycle.

To conclude, this study evidenced that ddPCR can be optimized to amplify targets coming from a complex matrix such as peat and can be used to target both universal markers (*16S rRNA* gene) and specific markers (e.g., *23S rRNA*, *cbbL* or *pufM* genes). Taken together, the results of this study support the effective use of ddPCR to analyze target gene concentrations in different peatland types and at different depths. These results further highlight a complicated picture in which major and emerging CFMs may all play a key role in peatland microbial CO_2_ fixation. This requires further support in the future to better understand peatland C cycle. Using a cutting-edge method such as ddPCR can therefore help to better understand the relevance and contribution of CFMs for the soil C cycle. Absolute quantification, which simplifies the quantification of functional genes, provides a better understanding of microbial ecology and its underlying processes within the ecosystems functioning.

## Acknowledgements

This work has been supported by the MIXOPEAT (Grant No. ANR-17-CE01-0007 to VEJJ) and BALANCE (Grant No. ANR-23-ERCC-0001-01) projects funded by the French National Research Agency. This study has been partially supported through the grant EUR TESS N°ANR-18-EURE-0018 in the framework of the Programme des Investissements d’Avenir.

