## Supplementary material for "Development of a ddPCR approach for the absolute quantification of soil microorganisms involved in atmospheric CO_2_ fixation"

#### Development of a ddPCR approach for the absolute quantification of microorganisms involved in peatlands carbon cycle

Included files:

- 7 supplementary figures
- 5 supplementary tables

### Supplementary Figures

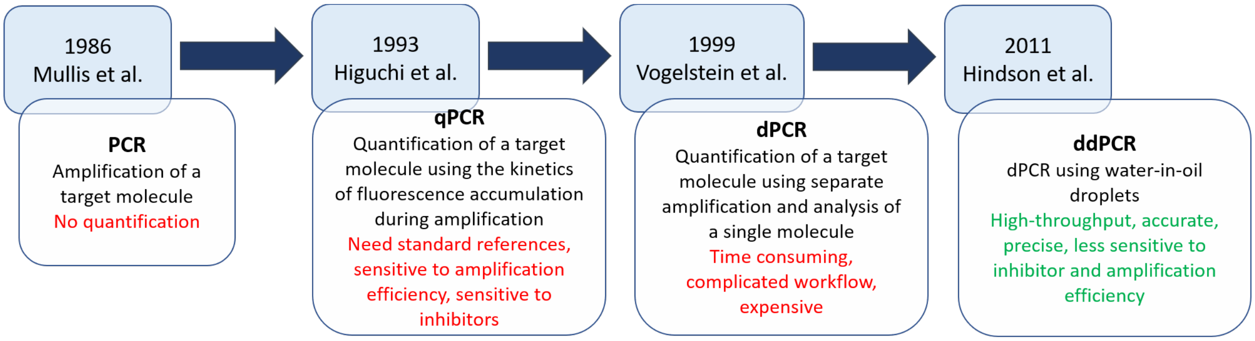


**Figure S1. Different molecular methods to quantify DNA** (inspired from Hou et al., 2023).

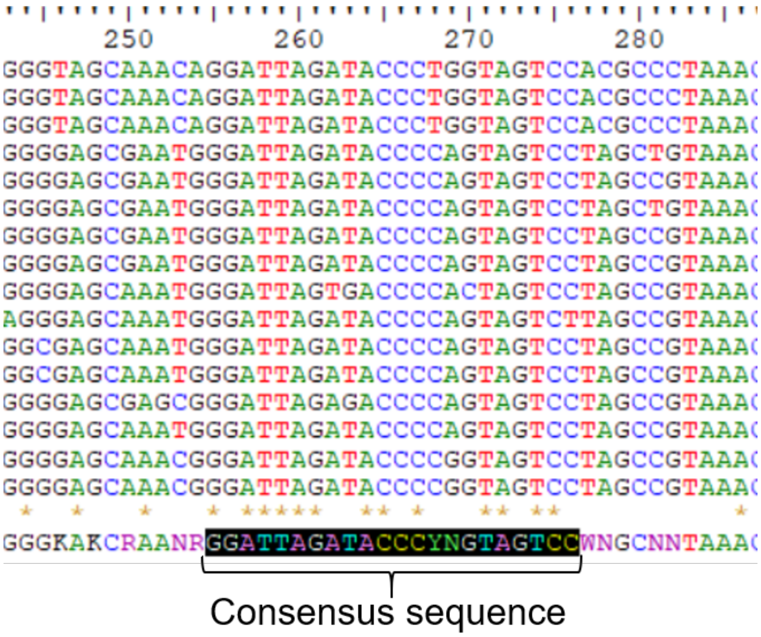


**Figure S2. Aligned sequence and consensus sequence for the design of the 23S255f forward primer to target the *23S rRNA* gene with ddPCR.**

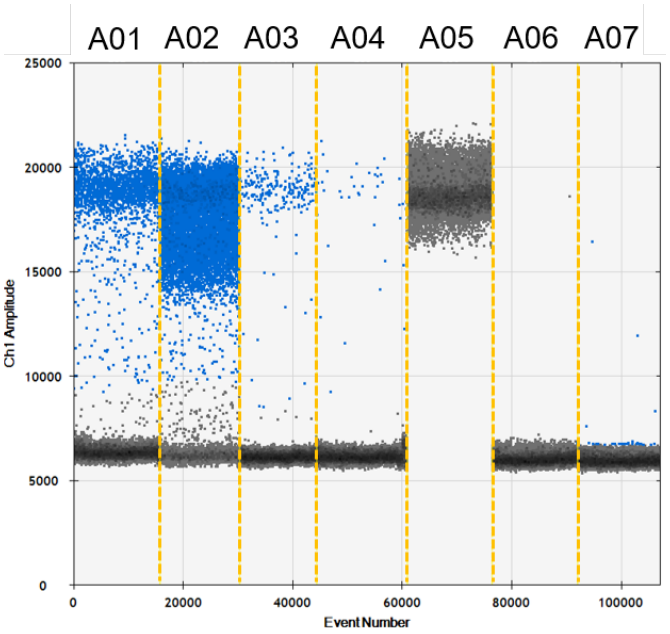


**Figure S3. Results of ddPCR assays for the *16S rRNA* gene targeting prokaryotes.** Only samples from the living layer (D1; 0-5 cm) from Counozouls were used for *16S rRNA* gene assays. A01 = DNA template diluted 10 times, A02 = DNA template not diluted, A03 = DNA template diluted 100 times, A04 = DNA template diluted 1,000 times, A05 = *E. coli* DNA (positive control), A06 and A07 = ultrapure water (negative controls).

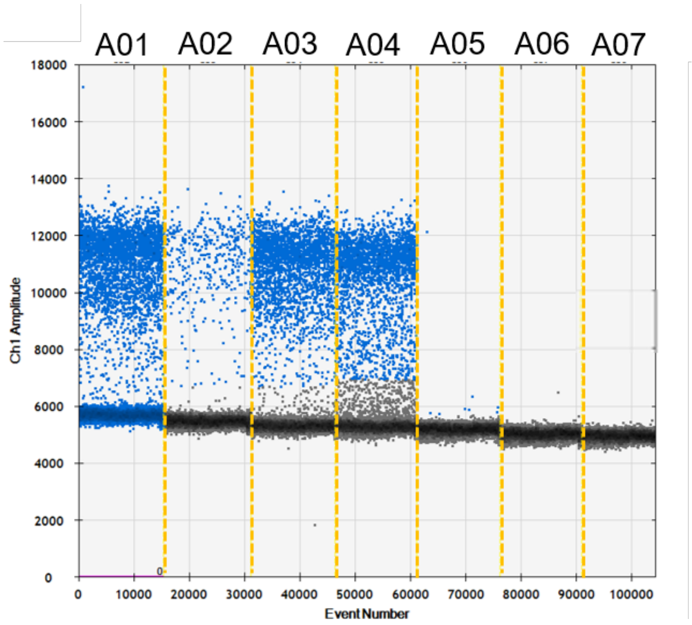


**Figure S4. Results of ddPCR assays for the *23S rRNA* gene targeting oxygenic phototrophs.** Samples from the living layer (D1; 0-5 cm), the decaying layer (D2; 5-10 cm) and the dead layer (D3; 10-15 cm) from Counozouls were used for *23S rRNA* gene assays. A01 = D1, DNA template diluted 10 times, A02 = D1, DNA template diluted 100 times, A03 = D2, DNA template diluted 10 times, A04 = D3, DNA template diluted 10 times, A05 = *E. coli* DNA (negative control), A06 and A07 = ultrapure water (negative controls).

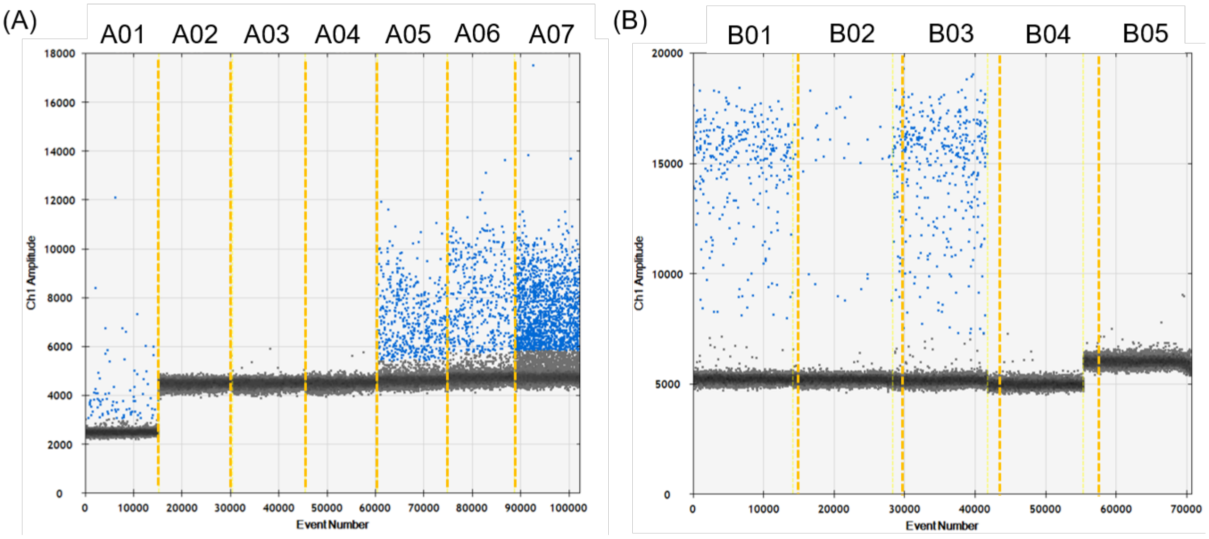


**Figure S5. Results of ddPCR assays for the *cbbL* gene targeting chemoautotrophs before (A) and after (B) optimization of ddPCR parameters.** Samples from the living layer (D1; 0-5 cm), the decaying layer (D2; 5-10 cm) and the dead layer (D3; 10-15 cm) from Counozouls were used for *cbbL* gene assays. Before ddPCR parameters optimization: A01 = D1, DNA template diluted 10 times, A02 and A03 = ultrapure water (negative controls), A04 = *E. coli* DNA (negative control), A05 = D3, DNA template diluted 10 times, A06 = D2, DNA template diluted 10 times and A07 = D1, DNA template not diluted. After ddPCR parameters optimization: B01 = D2, DNA template diluted 100 times, B02 = D2, DNA template diluted 1,000 times, B03 = D3, DNA template diluted 100 times, B04 = *E. coli* DNA (negative control) and B05 = ultrapure water (negative control).

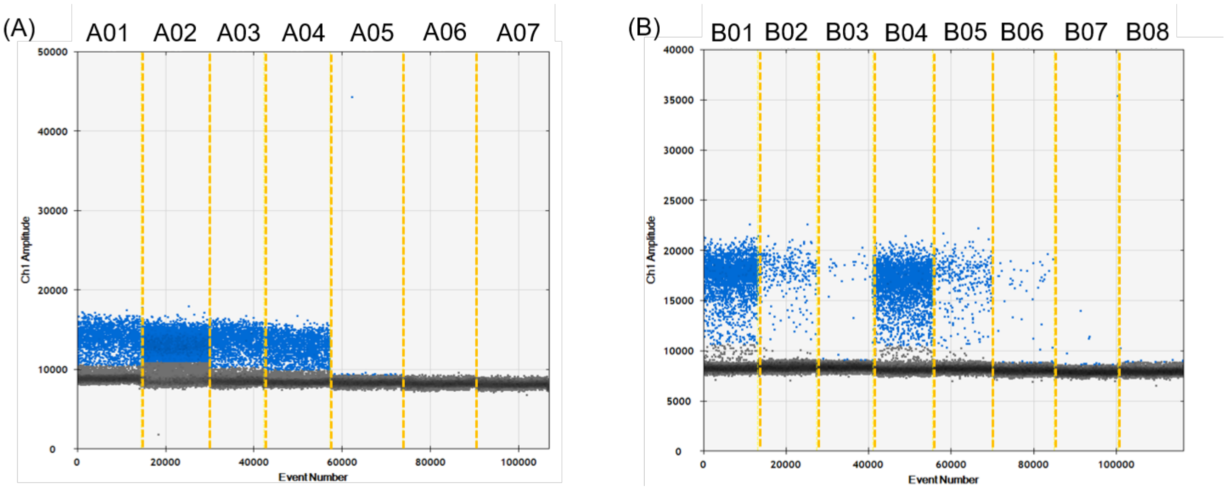

**Figure S6. Results of ddPCR assays for the *pufM* gene targeting AAnPBs before (A) and after (B) optimization of ddPCR parameters.** Samples from the living layer (D1; 0-5 cm), the decaying layer (D2; 5-10 cm) and the dead layer (D3; 10-15 cm) from Counozouls were used for *pufM* gene assays. Before ddPCR parameters optimization: A01 = D1, DNA template diluted 10 times, A02 = D1, DNA template not diluted, A03 = D2, DNA template diluted 10 times, A04 = D3, DNA template diluted 10 times, A05 = *E. coli* DNA (negative control), A06 and A07 = ultrapure water (negative controls). After ddPCR parameters optimization: B01 = D2, DNA template diluted 10 times, B02 = D2, DNA template diluted 100 times, B03 = D2, DNA template diluted 1,000 times, B04 = D3, DNA template diluted 10 times, B05 = D3, DNA template diluted 100 times, B06 = D3, DNA template diluted 1,000 times, B07 = *E. coli* DNA (negative control) and B08 = ultrapure water (negative control).

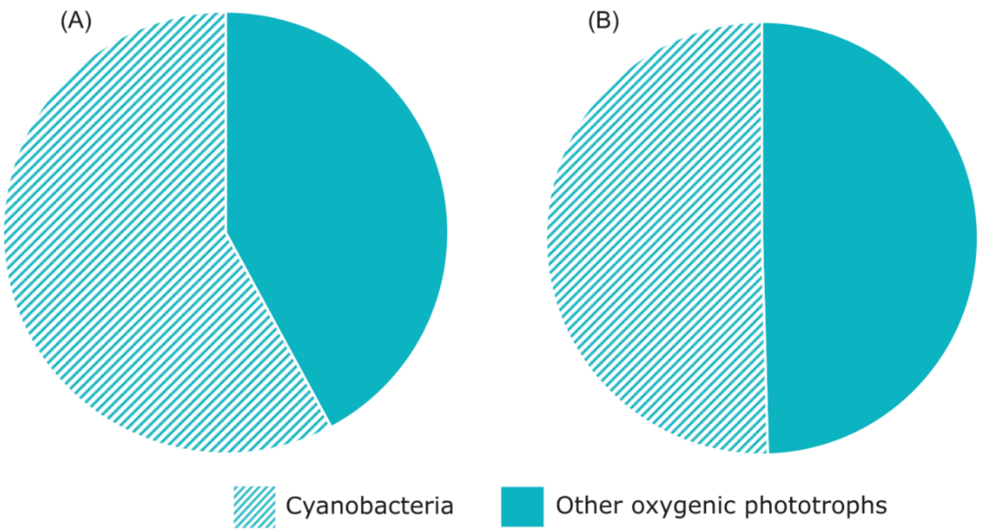


**Figure S7. Proportion of cyanobacteria among oxygenic phototrophs in Counozouls (A) and Männikjärve (B).**

### Supplementary Tables

**Table S1. Primers targeting the *23S rRNA* gene characteristics.**

| Primers | Forward | Reverse |
|---|---|---|
| Sequence | 5'- ggattagatacccydgtagtcc -3' | 5'- cagcctgttatccctagag -3' |
| Tm (°C) | 59.2 | 57.0 |
| GC (%) | 50 | 52.6 |
| Nucleotides | 22 | 19 |
| Degeneracy | 6 | 0 |
| Repetition of Gs or Cs longer than 3 bases | No | No |
| Self-dimer | No | Yes |
| Cross dimer | No | |
| Amplicon length | ~160 pb | |
| Ref | This study | (Sherwood & Presting, 2007) |

**Table S2. Results of ANOVA and post-hoc test for comparison of *16S rRNA* gene concentration between depths in Counozouls and in Männikjärve.**

| Depth | Counozouls | Männikjärve |
|---|---|---|
| D1 – D2 | $P$ = 0.029 | $P$ = 0.942 |
| D1 – D3 | $P = 6.5 \times 10^{-5}$ | $P$ = 0.176 |
| D2 – D3 | $P$ = 0.0082 | $P$ = 0.287 |

**Table S3. Results of ANOVA and post-hoc test for comparison of *23S rRNA* gene concentration between depths in Counozouls and in Männikjärve.**

| Depth | Counozouls | Männikjärve |
|---|---|---|
| D1 – D2 | $P$ = 0.189 | $P = 3.3 \times 10^{-4}$ |
| D1 – D3 | $P$ = 0.624 | $P = 4 \times 10^{-7}$ |
| D2 – D3 | $P$ = 0.038 | $P = 4.9 \times 10^{-4}$ |

**Table S4. Results of ANOVA and post-hoc test for comparison of *cbbL* gene concentration between depths in Counozouls and in Männikjärve.**

| Depth | Counozouls | Männikjärve |
|---|---|---|
| D1 – D2 | $P$ = 0.042 | $P$ = 0.789 |
| D1 – D3 | $P$ = 0.0014 | $P = 2.1 \times 10^{-4}$ |
| D2 – D3 | $P$ = 0.175 | $P = 6.1 \times 10^{-4}$ |

**Table S5. Results of ANOVA and post-hoc test for comparison of *pufM* gene concentration between depths in Counozouls and in Männikjärve.**

| Depth | Counozouls | Männikjärve |
|---|---|---|
| D1 – D2 | $P$ = 0.415 | $P$ = 0.065 |
| D1 – D3 | $P$ = 0.063 | $P = 3.1 \times 10^{-5}$ |
| D2 – D3 | $P$ = 0.463 | $P$ = 0.0015 |
